# Individual differences in functional brain connectivity predict temporal discounting preference in the transition to adolescence

**DOI:** 10.1101/255679

**Authors:** Jeya Anandakumar, Kathryn L. Mills, Eric Earl, Lourdes Irwin, Oscar Miranda-Dominguez, Damion V. Demeter, Alexandra Walton-Weston, Sarah Karalunas, Joel Nigg, Damien A. Fair

**Affiliations:** Department of Behavioral Neuroscience, Oregon Health & Science University, Portland, OR; Department of Psychology, University of Oregon, Eugene, OR; Department of Psychiatry, Oregon Health & Science University, Portland, OR; Advanced Imaging Research Center, Oregon Health & Science University, Portland, OR; Department of Psychology, University of Texas at Austin, Austin, TX

**Keywords:** delay discounting, fMRI, intrinsic connectivity, longitudinal, resting state

## Abstract

The transition from childhood to adolescence is marked by distinct changes in behavior, including how one values waiting for a large reward compared to receiving an immediate, yet smaller, reward. While previous research has emphasized the relationship between this preference and age, it is also proposed that this behavior is related to circuitry between valuation and cognitive control systems. In this study, we examined how age and intrinsic functional connectivity strength within and between these neural systems relate to changes in discounting behavior across the transition into adolescence. We used mixed-effects modeling and linear regression to assess the contributions of age and connectivity strength in predicting discounting behavior. First, we identified relevant connections in a longitudinal sample of 64 individuals who completed MRI scans and behavioral assessments 2-3 times across ages 7-15 years (137 scans). We then repeated the analysis in a separate, cross-sectional, sample of 84 individuals (7-13 years). Both samples showed an age-related increase in preference for waiting for larger rewards. Connectivity strength within and between valuation and cognitive control systems accounted for further variance not explained by age. These results suggest that individual differences in functional neural organization can account for behavioral changes typically associated with age.

## Introduction

Temporal discounting (also known as inter-temporal choice or delay discounting) is the process of assessing the value of waiting for a future reward depending on the magnitude of the reward and the delayed time. Individuals vary in their temporal discounting behavior, with some having a stronger preference for taking a smaller immediate reward versus waiting for a larger reward, and vice versa (Sadaghiani & Kleinschmidt, 2013). Previous experimental studies suggest a positive relationship between chronological maturation (age) and the tendency to prefer waiting for the larger reward (de Water, Cillessen, & Scheres, 2014; Steinberg et al., 2009), although some studies have found evidence for a nonlinear relationship in the transition into adolescence (Scheres, Tontsch, Thoeny, & Sumiya, 2014). Interestingly, the development of temporal discounting with age may be a stable marker of liability for disinhibitory psychopathologies such as ADHD even when psychopathological symptoms change with age (Karalunas et al., 2017). It has been proposed that brain function and organization can explain individual differences in temporal discounting behavior (Christakou, Brammer, & Rubia, 2011; Hare, Hakimi, & Rangel, 2014; Li et al., 2013; Scheres, de Water, & Mies, 2013; van den Bos, Rodriguez, Schweitzer, & McClure, 2014). Therefore, in this study, we analyzed how chronological maturation interacts with functional brain organization to predict temporal discounting.

### Temporal discounting as a measure of decision-making preference

Tasks assessing temporal discounting behavior can be used to measure an individual’s preference for a smaller-sooner reward (SSR) in comparison to a larger-later reward (LLR) (Green, Myerson, & Mcfadden, 1997). These tasks typically require individuals to choose between two rewards that vary in both the reward size and the delay time required until the amount is acquired (Myerson & Green, 1995). For example, participants typically respond to several questions in the following format: “At the moment, what would you prefer?” Below the question two options are presented (e.g. “$7.00 now”, “$10 in 30 days”). The SSR and LLR vary in both delay interval and reward size over successive trials; this way, the subjective value of temporal reward can be measured. Individuals preferring the SSR are characterized to have steeper temporal discounting; conversely, individuals preferring the LLR are characterized to have less temporal discounting. One way to measure this subjective value of temporal reward is through the use of indifference points (the delay duration at which the magnitude of SSR equals the magnitude of LLR) (Richards, Zhang, Mitchell, & de Wit, 1999). The indifference points are useful in calculating a single index of discounting rate, and in determining the value of the delayed reward (Yi, Pitcock, Landes, & Bickel, 2010). Specifically, plotting the indifference points in a series yields a discount curve, which describes the rate at which the value of reward decreases as time is increased.

### Neural networks involved in temporal discounting

Previous studies have shown that cortico-striatal circuitry is greatly involved in decision-making processes (Haber & Knutson, 2009), including temporal discounting (Peters & Büchel, 2011). In the present study, we focus on two cortico-striatal systems (defined *a priori*) that have been consistently correlated with different outcomes of an individual’s preference and value (Peters & Büchel, 2011; van den Bos et al., 2014): a valuation system (amygdala, medial orbitofrontal cortex, posterior cingulate cortex, ventromedial prefrontal cortex, and ventral striatum) and a cognitive control system (ventral lateral prefrontal cortex, dorsal anterior cingulate cortex, dorsolateral prefrontal cortex, dorsal striatum, and inferior frontal cortex) (See Figure **1a**). We also assessed connectivity between these networks and the supplementary motor area and hippocampus, given their involvement in intertemporal choice behavior (Peters & Büchel, 2010; Scheres et al., 2013; van den Bos et al., 2014). Overall, it has been theorized that adults with high temporal discounting preference are more likely to show greater recruitment of the control network and less recruitment of the valuation network when choosing a LLR over a SSR (van den Bos & McClure, 2013; Volkow & Baler, 2015).

**Figure 1:**
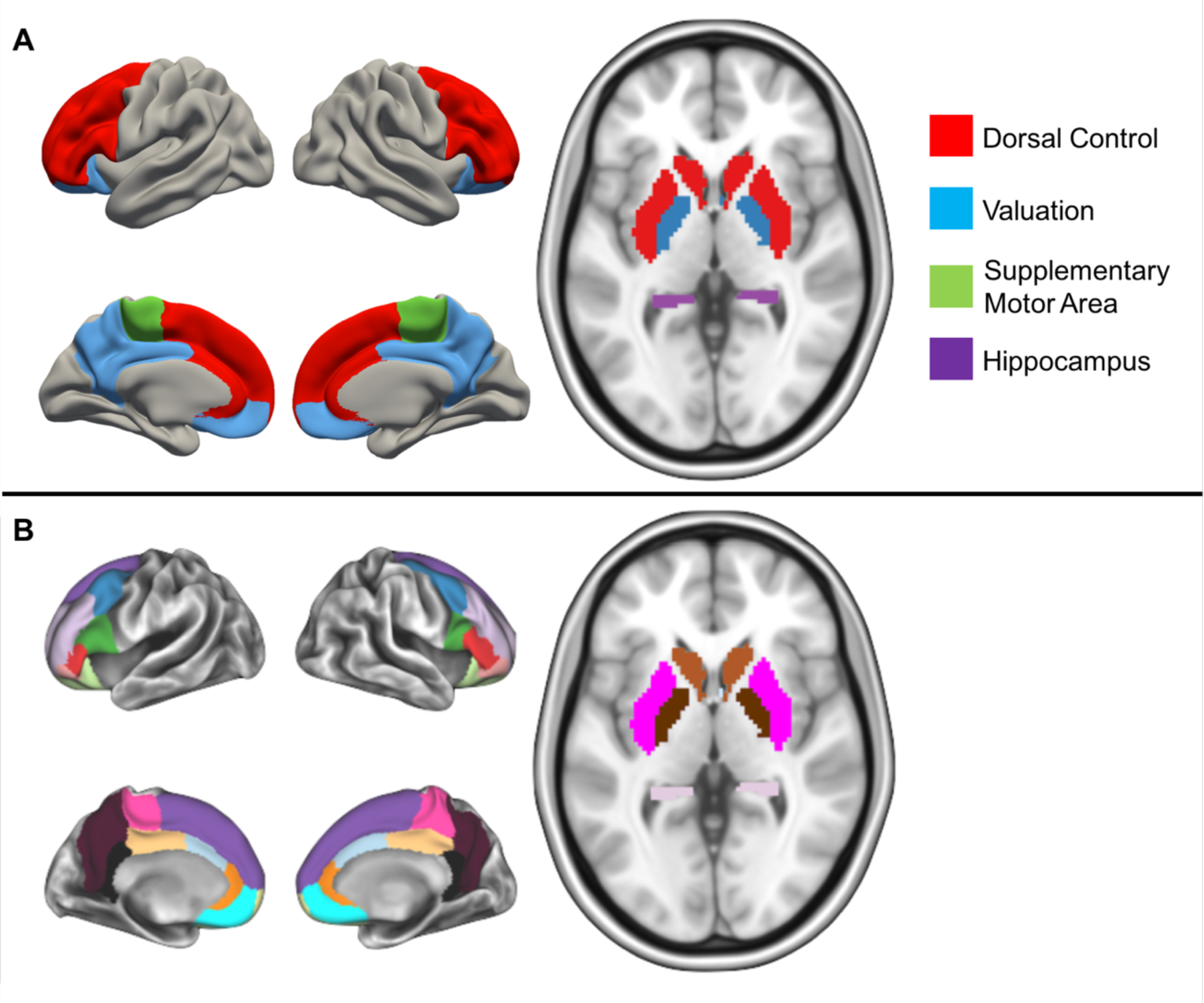
Brain systems of interest and regions of interest. [A] Brain networks (including two other regions out of the networks) included in this study. The regions in red represent the cognitive control network. The regions in blue represent the valuation network. The regions in green and purple represent the supplementary motor area and the hippocampus, respectively. [B] Each brain region included in this study.

Neural networks involved in temporal discounting can be interrogated with MRI in multiple ways, including task-based fMRI studies in which participants are asked to make temporal discounting decisions, and as well as in studies that compare anatomical or functional connectivity to temporal discounting preferences measured outside of the scanner. Previous studies have examined structural connectivity (white matter fiber integrity) and its relation to temporal discounting through Diffusion Tensor Imaging (DTI). Increased structural connectivity between the striatum and cortical control regions have been found to be related to decreased temporal discounting, whereas increased structural connectivity between the striatum and subcortical valuation regions were related to increased temporal discounting in adults (van den Bos et al., 2014).

While task-evoked brain activity can inform us on the functionality of cortical networks during specific contexts, intrinsic brain activity at rest can be used to measure an individual’s functional brain organization. The intrinsic activity of the brain reflects, in part, past activities, and these fluctuations impact future behavior (Sadaghiani & Kleinschmidt, 2013). Brain functionality and fluctuations are believed to determine and shape connectivity patterns. Here we study the brain’s intrinsic connectivity using resting-state functional connectivity MRI (rs-fcMRI) (Power, Schlaggar, & Petersen, 2014). rs-fcMRI measures the functional relationship between regions while the participant is not performing a specific task by measuring slow, spontaneous fluctuation of the blood oxygen level dependent (BOLD) signal. Intrinsic activity measures reveal the cohesive connections and interactions present in neuronal networks (Boly et al., 2008). Previous studies in adults have found that intrinsic brain connectivity within cortico-striatal networks were related to an individual’s temporal discounting preference (Calluso, Tosoni, Pezzulo, Spadone, & Committeri, 2015; Li et al., 2013).

### Development of neural networks underlying temporal discounting

It is hypothesized that differential rates of maturation across cortico-striatal systems, and the protracted development of the interconnections between them, are related to changes in behavior across development (Casey, 2015; Costa Dias et al., 2012, 2015; van den Bos, Rodriguez, Schweitzer, & McClure, 2015). In adults, it has been theorized that greater recruitment of control networks (and less recruitment of the valuation networks) are indicative of choosing the LLR, however, it is currently unclear if this brain-behavior relationship is present throughout development. One of the first task-based fMRI studies of temporal discounting examined the impact of age-related (ages 12-32 years; males) changes in brain activation when deciding between a SSR and a LLR (Christakou et al., 2011). This study demonstrated that when choosing an immediate reward, *increased* recruitment of the vmPFC and *decreased* recruitment of the ventral striatum, insula, anterior cingulate, occipital, and parietal cortices was related to increasing age and preference for LLR. Further, greater coupling between the ventral striatum and vmPFC was also related to increasing age and preference for LLR, suggesting that increased functional connectivity between the vmPFC and ventral striatum (regions of the valuation network) might be one neural mechanism underlying developmental changes in the preference for delayed rewards.

Another theory is that neural systems involved in three cognitive processes: valuation (i.e., the value placed on a certain stimuli or outcome), cognitive control (i.e., engaging in goal-directed cognitive processes), and prospection (i.e., thinking about the future), are involved in the process of temporal discounting (Peters & Büchel, 2011). Using this framework, Banich et al. (2013) compared the behavioral and neural correlates of temporal discounting in younger (14-15 years) and older (17-19 years) adolescents, and how these measures related to an individual’s self-reported tendency to think beyond the present. Behaviorally, older adolescents were more likely to choose a delayed reward over an immediate reward, and were slower than younger adolescents to choose the immediate reward (Banich et al., 2013). The pattern of brain activity related to intertemporal decision making was more distinct when choosing between immediate versus delayed rewards in the older adolescents compared to the younger adolescents (Banich et al., 2013). Across groups, individuals who reported a greater tendency to think beyond the present showed decreased recruitment of cognitive control regions during the temporal discounting task. These results suggest that both age and individual differences are related to the neural processing of temporal discounting.

Another study found that greater white matter integrity in pathways connecting the frontal and temporal cortices with other areas of the brain were positively correlated with the preference for delayed rewards across ages 9-23 years (Olson et al., 2009). Some of these correlations were developmentally related, whereas some of the effects appeared to be age-independent. For example, the relationship between greater white matter integrity in right frontal and left temporal regions and increased preference for delayed reward was not attributable to age. However, the relationship between integrity of white matter in left frontal, right temporal, right parietal (as well as some subcortical-cortical circuits) and the preference for delayed reward was age-related, as these white matter tracts also increased in integrity across the age range studied. These results show that both age and individual differences in neural circuitry are related to an individual’s preference for immediate versus delayed rewards. Another study examined the relationship between temporal discounting and fronto-striatal circuitry in a longitudinal study of individuals between the ages of 8-26 (Achterberg, Peper, van Duijvenvoorde, Mandl, & Crone, 2016). This study found that preference for LLR increased non-linearly between childhood and early adulthood, and that greater fronto-striatal white matter integrity was related to the preference for LLR (Achterberg et al., 2016).

Taken together, these studies demonstrate that people, on average, show increasing preference to wait for larger rewards rather than take immediate (smaller) rewards as they get older, but the increase may be nonlinear. Individual differences across development in temporal discounting preference are related to differences in functional neural organization. How one comes to choose a smaller immediate reward over a larger distant reward could be related to how that individual values the proposed reward, or it could be related to how well that individual can inhibit reflexive urges or the ability to think about the future. The development of brain systems involved in evaluating rewards, cognitive control, and thinking about the future all appear to contribute to the developmental changes in how we process situations that involve us making a choice between an immediate outcome and a distant outcome.

### Current study

This current project examines how developmental changes in functional connectivity between and within the cognitive control network, valuation network, hippocampus and SMA relate to temporal discounting preferences during the transition into adolescence. Specifically, we tested to see if changes in functional connectivity strength could explain additional variance in temporal discounting preferences above chronological age. Previous studies have reported no significant difference in discounting behavior between boys and girls (Cross, Copping, & Campbell, 2011; Lee et al., 2013), suggesting any sex effects are likely to be small. Therefore, to conserve statistical power, the relationship between sex and temporal discounting behavior was not examined.

## Methods

### Participants

Our two neurotypical samples were drawn from an ongoing longitudinal project examining brain development in children, recruited from the community, with and without attention-deficit/hyperactivity disorder (ADHD). Our first sample consisted of 64 individuals with 2 or 3 longitudinal scans each (n=137 scans) and our second cross-sectional sample consisted of 84 individuals. Details for both samples are included in **Table 1**. All participants were typically-developing children without psychiatric diagnoses and exhibited typical neurological patterns of thoughts and behavior throughout the study. Psychiatric status was evaluated based on evaluations with the Kiddie Schedule for Affective Disorders and Schizophrenia (KSADS; Puig-Antich & Ryan, 1986) administered to a parent; parent and teacher Conners’ Rating Scale-3rd Edition (Conners, 2003); and a chart review a child psychiatrist and neuropsychologist that required agreement. Any participant who was identified as having a current psychiatric, neurological, or neurodevelomental disorder was excluded from the present study. IQ was estimated with a three-subtest short form (block design, vocabulary, and information) of the Wechsler Intelligence Scale for Children, 4th Edition (Wechsler, 2003).

**Table 1.** Participant demographic characteristics for each sample.

|  | Longitudinal Sample Characteristics |  |  |  | Cross-sectional Sample Characteristics |  |  |
| --- | --- | --- | --- | --- | --- | --- | --- |
|  | All | Female | Male |  | All | Female | Male |
| N | 64 | 23 | 41 |  | 84 | 42 | 42 |
| Age mean (SD) | 10.8 (1.83) | 10.6 (1.95) | 10.9 (1.77) |  | 10.3 (1.39) | 10.3 (1.34) | 10.3 (1.44) |
| Age range | 7.3-15.7 | 7.3-15.7 | 7.5-14.5 |  | 7.3-13.3 | 8-13.3 | 7.2-13.2 |
| AUC mean (SD) | 0.51 (0.273) | 0.51 (0.261) | 0.51 (0.281) |  | 0.45 (0.288) | 0.44 (0.306) | 0.47 (0.273) |
| AUC range | 0.04 - 1 | 0.07 - 0.99 | 0.04 - 1 |  | 0.02 - 0.98 | 0.03 - 0.98 | 0.02 - 0.98 |
| IQ mean (SD) | 115.3 (13.95) | 116.6 (9.58) | 114.6 (15.88) |  | 116.5 (13.82) | 114.5 (14.86) | 118.4 (12.59) |
| IQ range | 72 - 144 | 98 - 132 | 72 - 144 |  | 78 - 148 | 78 - 144 | 96 - 148 |
| N visits | 137 | 49 | 88 |  | 84 | 42 | 42 |
| 2 visits | 55 | 20 | 35 |  | - | - | - |
| 3 visits | 9 | 3 | 6 |  | - | - | - |

### Temporal Discounting task

The temporal discounting task evaluates personal preference for a hypothetical delayed or immediate reward. Participants were presented a computerized task with a series of questions, and were read the following instruction before proceeding to the task:

> For the next task, you can choose between two options by clicking on it using the computer mouse. You can change your selection as often as you would like. Once you have decided which option you prefer, you can go on to the next question by clicking on the ‘next question’ box. One option will always be some amount of money available now. The other option will always be some amount of money later. The waiting period will vary between now and 180 days. Imagine that the choices you make are real– that if you choose ‘money now’ you would receive that amount of money at the end of the task and that if you choose ‘money later’ that you would actually have to wait before receiving the money. So, what are you going to do?

The computer-based task consisted of 92 questions with an option to get a reward immediately or get a larger amount of money ($10.00) at a later time period. Most of the participants were presented delays in intervals of 7, 30, 90, 180 days; a small percent of the participant were presented with different delay intervals of 1, 7, 30, 90 days.

Our temporal discounting task was analyzed by multivariate mathematical equations to measure an individual’s decision-making preference. Reward in relation to the time span is usually used to measure the preference of an individual or a collective population generalized by age. There are many mathematical ways to analyze temporal discounting task, however, for this experiment we choose Area Under Curve (AUC). AUC (see **Box 1**) best represents the preference of the participants as it takes into consideration the indifference point and the corresponding delay time (Myerson, Green, & Warusawitharana, 2001). AUC is equated to best represent the variables present in this experiment; it takes into account the sum of indifference and delay points acquired through temporal discounting, and outputs one value making it easier for analysis (Myerson et al., 2001).

#### Box 1.

##### AUC Equation

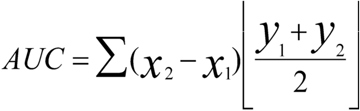

The *x*_2_ and *x*_1_ are the delayed points, and *y*_2_ and *y*_1_ represent the indifference points that correspond to the delays (Hamilton et al., 2015; Odum, 2011). The AUC outputs a signal value between 0 and 1; the lower the number represents the greater possibility to disregard the value of the reward, and have less tolerance for the delay time (Myerson et al., 2001; Odum, 2011). The AUC values and temporal discounting are inversely proportional, the closer the AUC value is to zero the more temporal discounting is present, therefore the participant is least likely to wait for a bigger reward. Likewise, the farther away the AUC value is to zero the most likely the participant is going to wait for the larger reward to be received at a later time.

Three validity criteria were applied to the quantification of AUC. The first criterion was to make sure that an indifference point for a specific delay was not greater than the preceding delay indifference point by more than 20% or $2 (Johnson & Bickel, 2008). The next criterion was the requirement for the final indifference point, at 180 days, to be less than the first indifference point, at 0 days, to indicate variation in subjective value of rewards across (Johnson & Bickel, 2008). The final criterion was to require the first, at 0 day, indifference point to be at least 9.25. This last criterion was enforced because a lower value indicates that the participant chose multiple time to receive the smaller “now” over the larger “now”, suggesting poor task engagement of misunderstanding of the task (Mitchell, Wilson, & Karalunas, 2015).

### MRI acquisition

MRI was acquired using a 3.0 Tesla Siemens Magnetom Tim Trio scanner with a twelve-channel head-coil at the Oregon Health & Science University Advanced Imaging Research Center. One high-resolution T1-weighted MPRAGE (TR=2300ms, TE=4ms, FOV=240×256mm, 1mm isotropic, sagittal acquisition) and multiple T2-weighted echo planar imaging (TR=2500ms, TE=30ms, FOV = 240×240mm, 3.8mm isotropic, either 82 or 120 volumes, axial acquisition, 90° flip angle) series were acquired during each scan visit. Functional data were collected at rest, in an oblique plane (parallel to anterior commissure-posterior commissure plane), and steady state magnetization was assumed after five frames (-10s). Participants were instructed to stay still and fixate their gaze on a standard fixation-cross in the center of the display during the acquisition of resting state scans.

### Image processing

The data were processed following the minimum processing steps outlined by the Human Connectome Project (Glasser et al., 2013), which included the use of FSL (Jenkinson, Beckmann, Behrens, Woolrich, & Smith, 2012; Smith et al., 2004; Woolrich et al., 2009) and FreeSurfer image analysis suite (http://surfer.nmr.mgh.harvard.edu/) (Dale, Fischl, & Sereno, 1999; Fischl, Sereno, & Dale, 1999). With this method, gradient distortion corrected T1w and T2w volumes are first aligned to MNI’s AC-PC axis and then nonlinearly normalized to the MNI atlas. Next, the T1w and T2w volumes are re-registered using boundary based registration (Greve & Fischl, 2009) to improve alignment. The brain of each individual is then segmented using the ‘recon-all’ FreeSurfer functions, which are further improved by utilizing the enhanced white matter-pial surface contrast of the T2w sequence. The initial pial surface is calculated by finding voxels that are beyond ± 4 standard deviations from the grey matter mean. The resulting parameter is then used to make sure no lightly myelinated grey matter is excluded. The estimated segmentation is refined further by eroding it with the T2w volume. Of the 221 total scan visits included in this study, 51 (23%) were processed without a T2w volume, either because this sequence was not acquired or was judged as being of low quality. These 51 were processed using FreeSurfer’s regular T1 segmentation algorithm (Fischl et al., 2002). Next, the preliminary pial surface and white matter surface are used to define an initial cortical ribbon. The original T1w volume is smoothed with the ribbon using a Gaussian filter with a sigma of 5mm. Then, the original T1w image is divided by the smoothed volume to account for low frequency spatial noise. This filtered volume is used to recalculate the pial surface, but now using 2 (instead of 4) standard deviations as threshold to define the pial surface. These segmentations are then used to generate an individualized 3D surface rendering of each individual, which is finally registered to the Conte 69 surface atlas as defined by the Human Connectome Project. This registration process allows all data types (cortical thickness, grey matter myelin content, sulcal depth, function activity, functional and structural, connectivity, etc.) to be aligned directly within and between individuals. All T1w and T2w MRI scans were quality controlled for any noticeable movement through visual inspection of raw and reconstructed images. The images were assessed in a pass or fail manner; scans that failed were excluded from the samples included in the present study.

Functional EPI data are registered to the first volume using a 6-degrees of freedom linear registration and correcting for field distortions (using FSL’s TOPUP), except for two scans (of 221) where no field map had been acquired. Next the EPI volumes are averaged, with each volume of the original time series re-registered to the average EPI volume using a 6-degrees of freedom linear registration. This last step avoids biases due to a single frame being used, which may be confounded by variability of movement across a given run. The average EPI volume is then registered to the T1w volume. The matrices from each registration step are then combined, such that each frame can be registered to the atlas all in a single transform (i.e. only one interpolation).

The resulting time-courses are then constrained by the grey matter segmentations and mapped into a standard space of 91,282 surface anchor points (greyordinates). This process accounts for potential partial voluming by limiting the influence of voxels that “straddle” grey and non-grey matter voxels (pial surface, white matter, ventricles, vessels, etc). Two thirds of the greyordinates are vertices (located in the cortical ribbon) while the remaining greyordinates are voxels within subcortical structures. Thus, the BOLD time courses in greyordinate space are the weighted average of the volume’s time courses in grey matter, where the weights are determined by the average number of voxels wholly or partially within the grey matter ribbon. Voxels with a high coefficient of variation are excluded. Next, the surface time courses are downsampled to the greyordinate space after smoothing them with a 2mm full-width-half-max Gaussian filter.

The additional preprocessing steps necessary for resting-state functional connectivity analyses consist of regressing out the whole brain (in this case the average signal across all greyordinates (e.g., see Burgess et al., 2016), ventricle and white matter average signal, and displacement on the 6 motion parameters, their derivatives and their squares (Power, Mitra, et al., 2014). All regressors are individualized and specific to the participant, based on their own segmentations. The regression’s coefficients (beta weights) are calculated solely on the frames where the frame displacement is below 0.3mm to reduce the influence of movement “outliers” on the output data, but all the time courses are regressed to preserve temporal order for temporal filtering. Finally, time courses are filtered using a first order Butterworth band pass filter with cutting frequencies of 9 millihertz and 80 millihertz.

We applied a strict motion censoring procedure to the resting-state images (Fair, Nigg, et al., 2012; Power, Barnes, Snyder, Schlaggar, & Petersen, 2012) which takes the absolute value of the backward-difference for all rotation and translation measures in millimeters, assuming a brain radius of 50mm, and summates those absolute backward-differences for a measure of overall framewise displacement (FD). Volumes with a displacement exceeding 0.2mm were excluded, and we also removed frames with less than five contiguous frames of low motion data between instances of high motion (FD > 0.2mm) data to confidently account for motion effects on adjacent volumes (Power, Mitra, et al., 2014). Only participants with greater than 5 minutes of high quality data were included in the present analysis. The mean framewise displacement of participants in the first sample was 0.08 ± 0.02mm; range 0.05 - 0.13mm. The mean framewise displacement of participants in the second sample was 0.09 ± 0.02mm; range 0.04 - 0.13mm. More information on the motion characteristics on the full sample (i.e. including those excluded) can be viewed in Dosenbach et al., (2017).

### Regions of interest

Our regions of interest (ROIs) included regions within valuation and cognitive control systems, as well as hippocampus and supplementary motor area (SMA). For our cortical ROIs, we selected regions within each of these networks from the Deskan-Killiany atlas provided by FreeSurfer (Desikan et al., 2006). While other parcellations can be considered, we chose this parcellation in order to examine anatomically-defined cortical regions that have been identified in previous work. Cortical reconstruction and volumetric segmentation was performed with the FreeSurfer image analysis suite, which is documented and freely available for download online (http://surfer.nmr.mgh.harvard.edu/). The technical details of these procedures are described in prior publications (Dale et al., 1999; Fischl et al., 2002; Fischl & Dale, 2000). FreeSurfer uses individual cortical folding patterns to match cortical geometry across subjects (Fischl et al., 1999), and maps this parcellation of the cerebral cortex into units with respect to gyral and sulcal structure (Desikan et al., 2006; Fischl et al., 2004). Our striatal and subcortical ROIs were defined based on FreeSurfer’s anatomical segmentation procedure. For the purposes of this study we examined the nucleus accumbens (NAcc), pallidum, amygdala, medial orbitofrontal cortex (mOFC), and posterior cingulate cortex (PCC) as part of the valuation network, and the caudate, putamen, anterior cingulate cortex (ACC), dorsal anterior cingulate cortex (dACC), dorsolateral prefrontal cortex (dlPFC), inferior frontal gyrus (IFG), and ventrolateral prefrontal cortex (vlPFC) of the cognitive control network (**Figure 1; SI Table 1**).

### Statistical analysis

In this study, we first tested to see whether chronological age could be used to predict temporal discounting preference as measured by AUC. We then tested to see if the strength of connectivity between each of our ROIs was able to explain variance in temporal discounting AUC values above chronological age. All analyses were conducted in R version >3.3.3 (https://www.r-project.org/). The script we used to conduct these analyses is freely available online to facilitate reproducibility and replication efforts (https://github.com/katemills/temporal_discounting).

### Sample 1

For our first, longitudinal, sample we tested each of these questions using mixed-effects models with the nlme package implemented through R. Mixed effects modeling accounts for the non-independence of the data collected from the same individual over time, and allows for unequal spacing between data collection points. This statistical analysis contains both the average slope and intercepts of the parameter (fixed effects), and varying intercept for each individual that is a random deviation of the fixed effect (random effect). We tested the following three polynomial models to predict AUC from chronological age:

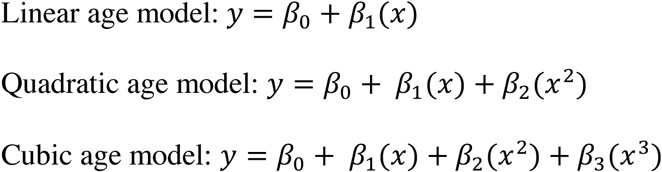

Where *y* is the AUC value, and *β*_0_ represents the intercept; “ represents the participant’s age; and *β_1_*, *β_2_* and *β_3_* represent regression coefficients. We centered age for all analyses (10.70 years). The three age models were compared and tested against a null model that only included the random intercept for each individual. The best fitting model was determined by Akaike Information Criterion (AIC) and likelihood ratio (LR) statistics using the heuristic of parsimony. The model with the lowest AIC value that was significantly different (*p*<.05), as determined from LR tests, from less complex models was chosen.

To identify the connections that could predict an individual’s AUC score above chronological age, we used LR statistics to compare models including a connection of interest (COI) correlation coefficient as an interaction and/or main effect added to the age only model. These brain connectivity models were then compared against each other as well as the best fitting age model. The model with the lowest AIC value that was significantly different (*p*<.01) from less complex models was selected as the best fitting model. To account for the possibility that brain connectivity alone could account for more variance in AUC values than the age-only model or the multivariate models, we also tested to see if a model including the COI correlation coefficient, but not age, was the best fitting model. We identified connectivity-only models if they had lower AIC than the age only models, and were also both significantly different and had lower AIC than the other more complex models.

### Sample 2

We examined the same questions in the second sample to test the replicability of the results obtained from the first sample. Similar to our first sample, we first examined the relationship between AUC and chronological age, specifically by comparing linear to nonlinear models (quadratic & cubic). Since these data were cross-sectional, we used regular linear regression to fit these models and compared models through F tests (*p*<.05). Age was centered for all analyses (10.23 years). Once the best age model was determined, we tested if adding COI correlation coefficients to the model would improve the model fit through F tests (*p*<.05). We only examined the COIs that were determined to explain additional variance in AUC above age in the first sample.

## Results

### AUC increases from late childhood into early adolescence

Model comparisons between the null, linear age, quadratic age, and cubic age models are presented in **Table 2**. Of the three age models tested, the quadratic model best represented the relationship between age and AUC in this longitudinal sample (LR quadratic model vs. null: 13.2, *p* < .002). The results of this model suggest that, on average, each yearly increase in age across this sample was associated with an increase of 0.04 AUC, with a negative rate of change (−0.01) (**Table 3; Figure 2**). These results should be interpreted from the predicted intercept at age 10.70 years (0.55). The graph illustrates a group-level increase in AUC until age ~11 years, but relative stability in AUC between ages 11-14 years.

**Table 2.** Comparison of polynomial age models for the longitudinal sample.

| **Longitudinal sample** |  |  |  |  |  |  |  |  |
| --- | --- | --- | --- | --- | --- | --- | --- | --- |
|  | Mode<br>1 | df | AIC | BIC | logLik | Test | L.Ratio | p-value |
| Null Model | 1 | 3 | 25.0 | 33.7 | -9.5 |  |  |  |
| Linear Age | 2 | 4 | 18.8 | 30.4 | -5.4 | 1 vs 2 | 8.2 | 0.0042 |
| Quadratic Age | 3 | 5 | 15.8 | 30.4 | -2.9 | 2 vs 3 | 5.0 | 0.0257 |
| Cubic Age | 4 | 6 | 15.4 | 32.9 | -1.7 | 3 vs 4 | 2.4 | 0.1226 |

**Table 3.** Fixed effects for best fitting (quadratic) age model predicting AUC for the longitudinal sample.

| **Longitudinal Sample** |  |  |  |  |  |
| --- | --- | --- | --- | --- | --- |
|  | Value | Std. Error | DF | t-value | p-value |
| Intercept | 0.55 | 0.03 | 71 | 17.0 | <0.0001 |
| Linear age | 0.04 | 0.01 | 71 | 3.2 | 0.0021 |
| Quadratic age | -0.01 | 0.01 | 71 | -2.2 | 0.0291 |

**Figure 2:**
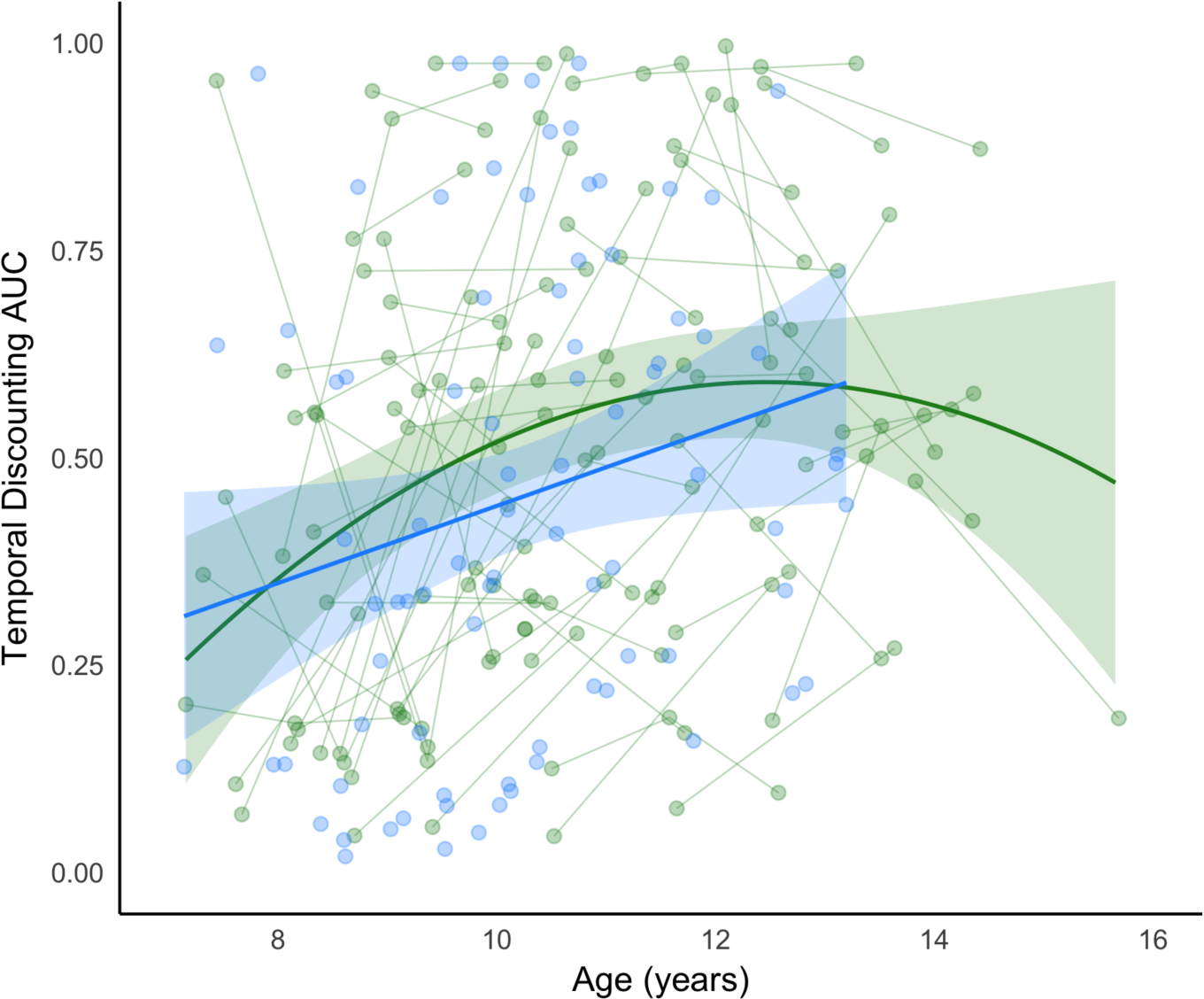
Best fitting age models for AUC. The green line represents the predicted model fit for AUC for sample 1 (longitudinal sample) and the blue line represents the predicted model fit for AUC for sample 2 (cross-sectional sample). Shading represents the 95% confidence intervals. Raw data are plotted in the background, with each individual measurement representing a circle, and lines connecting data collected from the same individual across time.

In our second, cross-sectional, sample, we found evidence for a linear relationship between age and AUC (**Figure 2**; blue). The linear model for this sample suggests that, on average, each yearly increase in age across this sample was associated with an increase of 0.05 AUC (**Table 3; Figure 2**). These results should be interpreted from the predicted intercept at age 10.23 years (0.45). Overall, the graph shows a similar increase in AUC across the age period studied as is visible in the longitudinal sample.

### Brain connectivity explains variance in AUC not accounted for by age

In the first sample, we found that AUC was best predicted by models including both age and connectivity for fifty-eight COIs **(SI Table 2)**. Many of the connections (40%) were between regions within the cognitive control network, whereas 10% of connections were between regions within the valuation network. 36% of the connections were between the cognitive control network regions and the valuation network regions. None of the identified connections included connections between the control network and the SMA or the hippocampus, however, one connection between the valuation network and hippocampus and three connections between the valuation and the SMA were identified as relevant to predicting AUC. All four possible connections between the SMA and hippocampus were identified as relevant to predicting AUC.

Of the fifty-eight connections identified in the first sample, only nine were replicated in the cross-sectional sample (**Table 4; Figure 3**). Three of the nine connections represented connections within regions of the cognitive control system (left dlPFC – right dACC; bilateral dlPFC; bilateral superior frontal cortex); three represented connections within regions of the valuation system (right pallidum – left PCC; right pallidum – right PCC; right mOFC – left amygdala); and three represented connections between these two systems (left dlPFC – right PCC; left superior frontal cortex – right PCC; left mOFC – right vlPFC). Model statistics for these nine models are detailed for both the longitudinal sample and cross-sectional sample in **Table 4**.

**Table 4.**
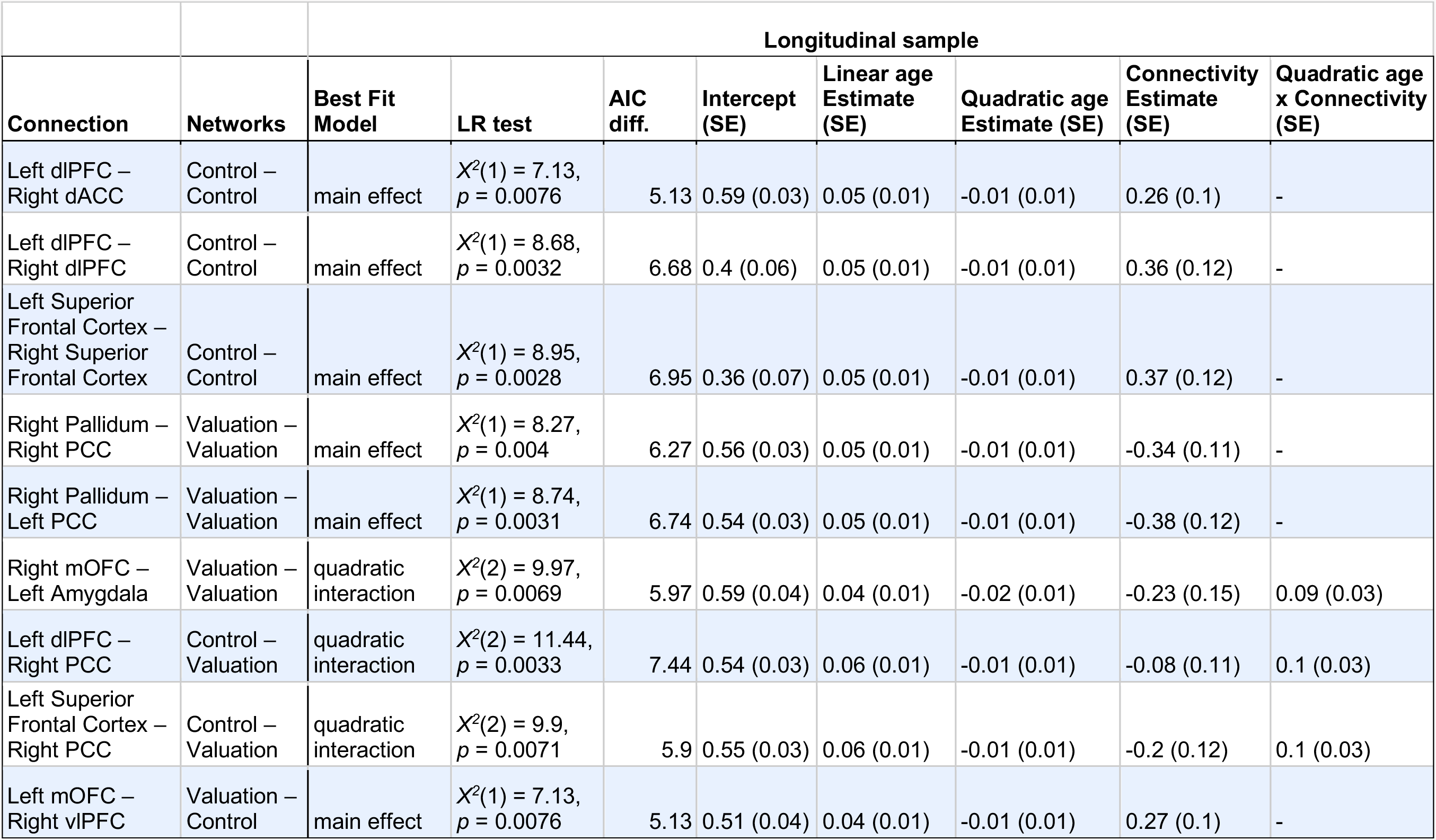
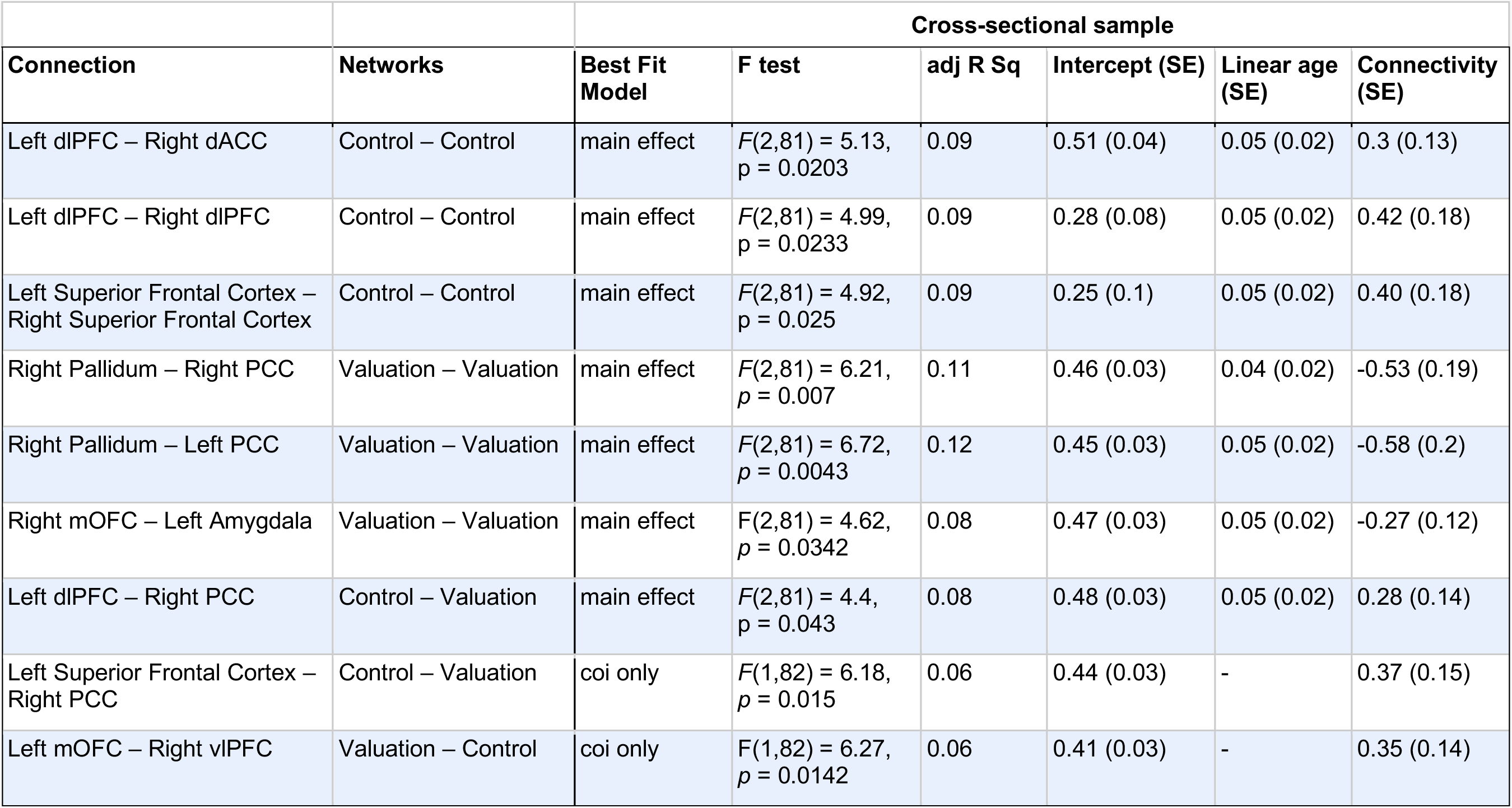
Best fitting model characteristics for the nine connections of interest that replicated across the both samples.

**Figure 3a-c:**
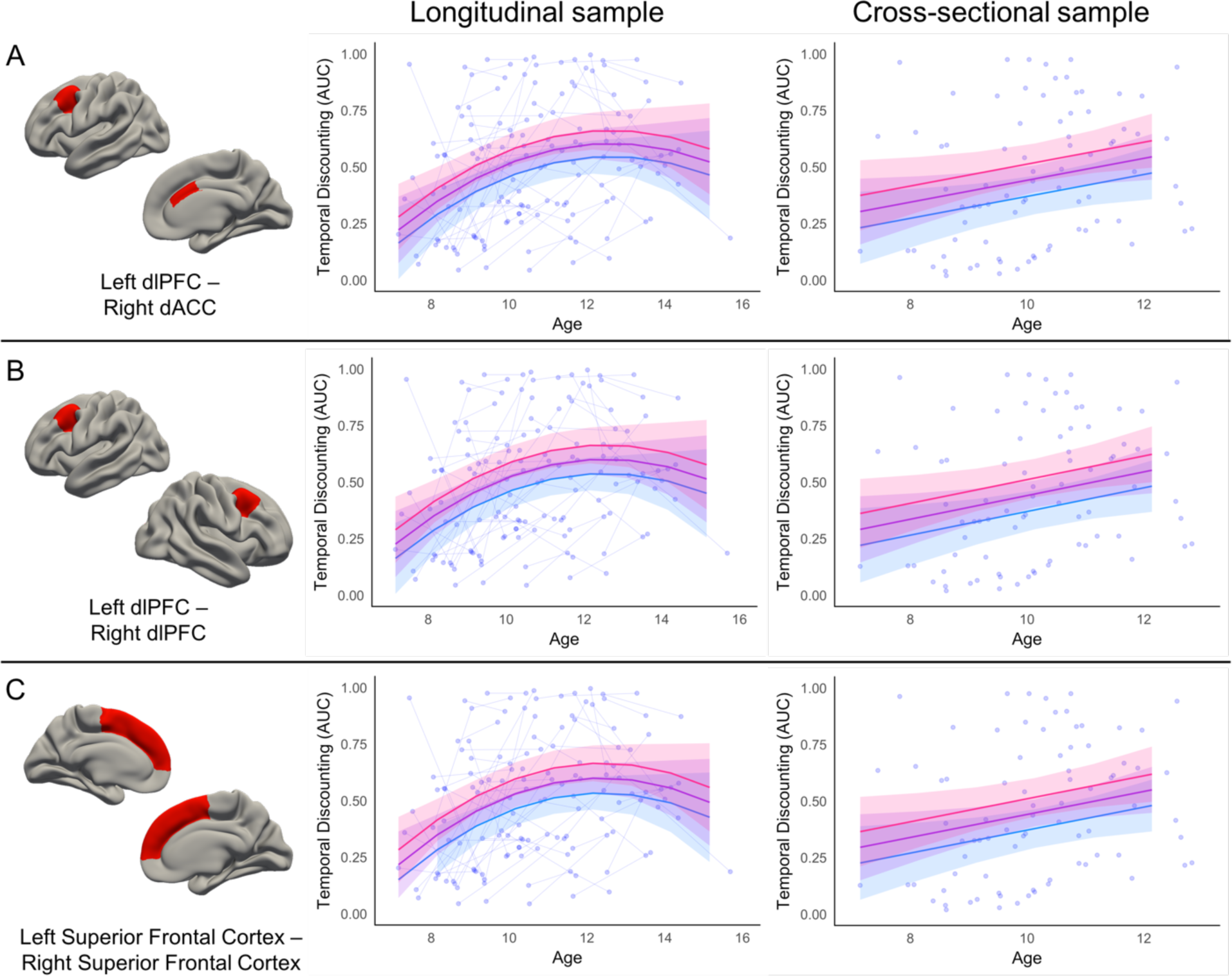
Relationship between cognitive control regions and AUC. The cortical regions involved in the connectivity between two cognitive control systems are represented by red on the brain. Pink trajectory represents AUC for an individual with 1 standard deviation higher connectivity than the mean between the two regions. Purple trajectory represents predicted AUC for participants with the mean connectivity strength between the two regions. Blue trajectory represents AUC for an individual with 1 standard deviation lower connectivity than the mean between the two regions. Raw data are plotted in the background, with each individual measurement representing a circle, and lines connecting data collected from the same individual across time.

The majority of the identified connections showed similar effects across samples. The three connections within the cognitive control system impacted the prediction of AUC similarly in both samples: individuals with greater connectivity strength between these cognitive control regions were predicted to have a preference for LLR (higher AUC) across the age ranges studied. The beta values for the main effect of connectivity were similar across the samples as well, with connectivity beta estimates ranging from 0.26 – 0.37 for the longitudinal sample, and 0.30 – 0.42 for the cross-sectional sample.

The three connections within the valuation system also impacted the prediction of AUC similarly in both samples: individuals with greater connectivity strength between these valuation regions were predicted to have a preference for the SSR (lower AUC) across the age ranges studied. The beta values for the main effect of connectivity were similar across the samples as well, with connectivity beta estimates ranging from −0.38 – −0.23 for the longitudinal sample, and −0.58 – −0.27 for the cross-sectional sample. The impact of connectivity between the right pallidum and PCC on predicting AUC with age was virtually identical for both cortical hemispheres.

Individuals with greater connectivity strength between the left mOFC and right vlPFC were predicted to have a preference for LLR (higher AUC) across the age ranges studied, similar to patterns found for connections between the cognitive control regions. Connectivity between these two regions was a better predictor of AUC than age alone in the cross-sectional sample. Within the longitudinal sample, connectivity strength between the right PCC and the left dlPFC or left superior frontal cortex interacted with the quadratic age term to predict AUC, with stronger connectivity strength predicting a preference for LLR (higher AUC) only at the tail ends of the age range. Within the cross-sectional sample, participants greater connectivity strength between the right PCC and left dlPFC were predicted to have a preference for LLR (higher AUC). Connectivity between the right PCC and left superior frontal cortex was a better predictor of AUC than age alone in the cross-sectional sample, with individual with greater connectivity strength between these regions having a preference for LLR (higher AUC).

## Discussion

In this study, we investigated whether individual differences in functional brain organization are associated with temporal discounting preferences in the transition into adolescence. Specifically, we tested if functional connectivity between regions involved in valuation, cognitive control, hippocampus and SMA could explain variance in temporal discounting preference (AUC) above chronological age. To ensure validity of our reported results, we tested these models in two independent datasets: a longitudinal dataset of children aged 7-15 years and a cross-sectional dataset of 7-13 year olds.

In both samples we observed a group-average increase in AUC between late childhood and early adolescence. We found evidence that the relationship between age and AUC was best represented by a quadratic trajectory in our longitudinal sample, with AUC increasing between ages 7-11 years before stabilizing. For the cross-sectional sample, we identified a linear increase in AUC between ages 7-13 years. While the best fitting model differed between these samples, the overall pattern observed in both samples reflected a general trend for our participants to prefer waiting for a later, larger reward (LLR) as they got older.

This result supports the general finding that temporal discounting preferences shift in the transition into adolescence (Achterberg et al., 2016; Scheres et al., 2014). Scheres et al., (2014) demonstrated in a cross-sectional sample encompassing ages 6-19 years that adolescents were more likely to wait for the LLR in comparison to children and young adults. Achterberg et al., (2016) similarly found that the ability to delay gratification increased from childhood into adolescence. It is important to note that, although we found a group-average increase in AUC across the transition into adolescence, there was substantial individual variability (see **Figure 2**). Further, because our sample age range ends at 15 years we cannot be sure if the preference for LLR declines between mid-to-late adolescence.

### Individual differences in functional connectivity are related to temporal discounting preference

The current study proposed that individual variability in temporal discounting preference could be explained by differences in intrinsic functional organization. To test this hypothesis, we examined if intrinsic functional connectivity between a set of *a priori* regions of interest and networks could improve the “age only” models in predicting an individual’s temporal discounting preference. To mitigate false positives and overfitting, we implemented both a stringent model selection procedure utilizing AIC as well as Likelihood Ratio tests paired with replication in an independent sample. We found nine distinct brain connections were able to explain variance in temporal discounting preference above age alone in both our longitudinal and cross-sectional samples. These findings suggest that individual differences in functional brain connectivity can explain a portion of individual variability in temporal discounting preferences during the transition to adolescence.

Our results demonstrate that individuals with greater connectivity between cortical regions within cognitive control systems are more inclined to choose LLR. Specifically, we found that increased connectivity between the left dlPFC and the right dACC, bilateral dlPFC, and bilateral superior frontal cortex, relate to a preference for LLR for individuals across the transition into adolescence (**Figure 3a-c**).

Across samples, we found evidence that greater connectivity between right pallidum and the bilateral PCC was associated with a preference for SSR across the transition into adolescence. Specifically, a greater connectivity between these valuation regions predicted lower AUC for individuals across the age ranges studied (**Figure 3de**). These results align with previous findings showing individual differences in cortico-striatal circuitry are related to temporal discounting preferences (van den Bos et al., 2014; 2015). Our results also demonstrate that increased connectivity between the left amygdala and right mOFC was related to increased preference for the SSR in the transition to adolescence (**Figure 3f**). While a main effect was found for the cross-sectional sample, there was an interaction between connectivity and the quadratic age term for the longitudinal sample. This presents the possibility that the relationship between increased connectivity between the left amygdala and right mOFC and temporal discounting preference is not static across ages 7-15 years.

**Figure 3d-f:**
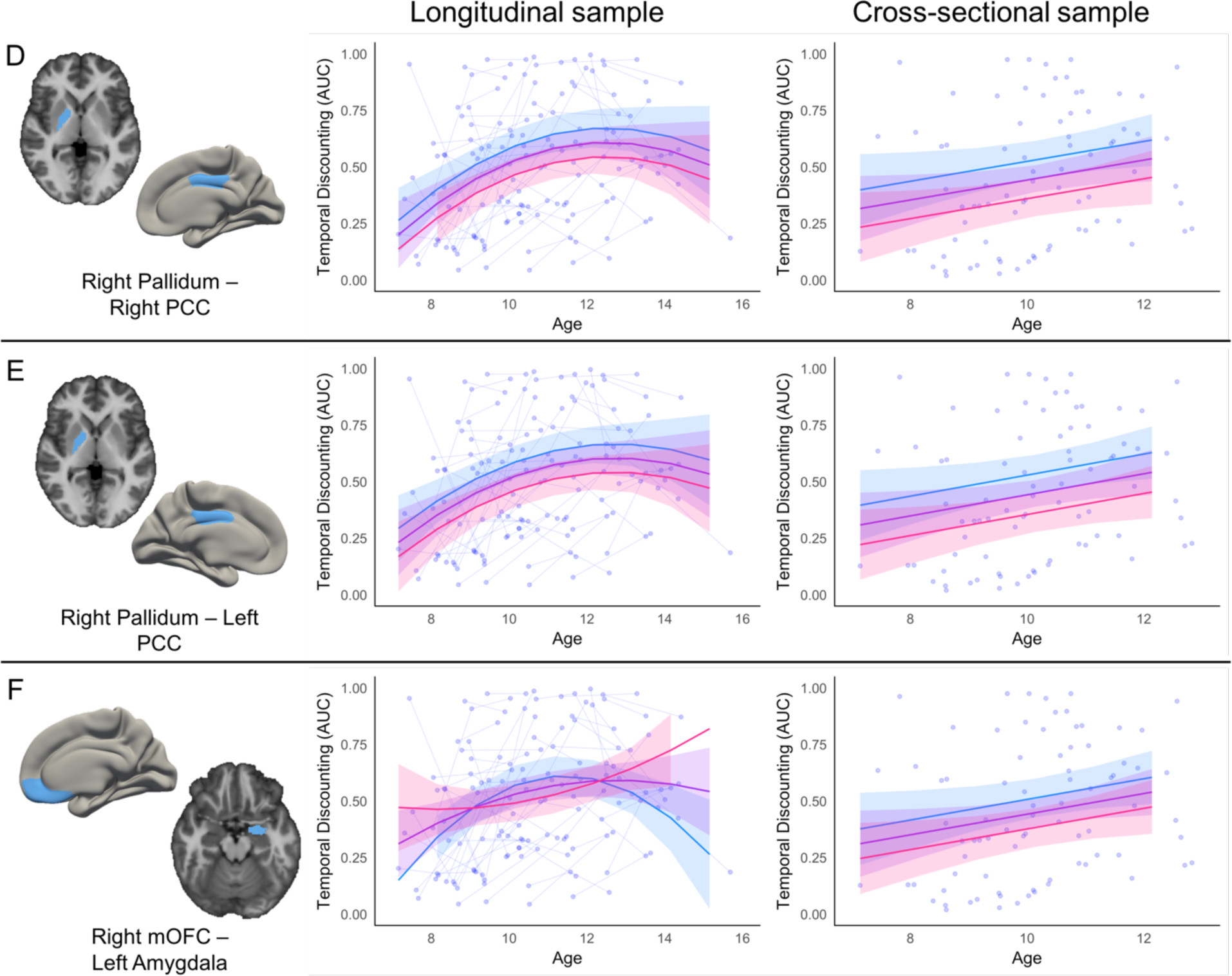
Relationship between valuation regions and AUC. The regions, between cortical and subcortical, involved in the connectivity between two valuation systems are represented by blue on the brain. Pink trajectory represents AUC for an individual with 1 standard deviation higher connectivity than the mean between the two regions. Purple trajectory represents predicted AUC for participants with the mean connectivity strength between the two regions. Blue trajectory represents AUC for an individual with 1 standard deviation lower connectivity than the mean between the two regions. Raw data are plotted in the background, with each individual measurement representing a circle, and lines connecting data collected from the same individual across time.

While we found evidence that increased connectivity between the right PCC and the left dlPFC or left superior frontal cortex was related to greater preference for LLR for individuals across ages in the cross-sectional sample, the best fitting models in the longitudinal sample suggested a nonlinear relationship between this strength of these connections and AUC preference across age (**Figure 3gh**). We found that greater connectivity between the left mOFC to right vlPFC (the pars orbitalis region of the inferior frontal gyrus) was related to increased preference for LLR across the transition into adolescence. This possibly reflects that stronger functional connectivity at rest between these regions reflects the ability for the vlPFC/IFG to regulate mOFC signaling (Hare, Camerer, & Rangel, 2009). In both samples, a main effect of greater connectivity between the dlPFC and several regions predicted higher AUC (increased preference for LLR or less discounting) for individuals across the transition into adolescence. This result held for connections between the dlPFC and dACC, bilateral dlPFC, as well as dlPFC and PCC, further underscoring the role of dlPFC in the development of temporal discounting behavior (Wang et al., 2017).

**Figure 3g-i:**
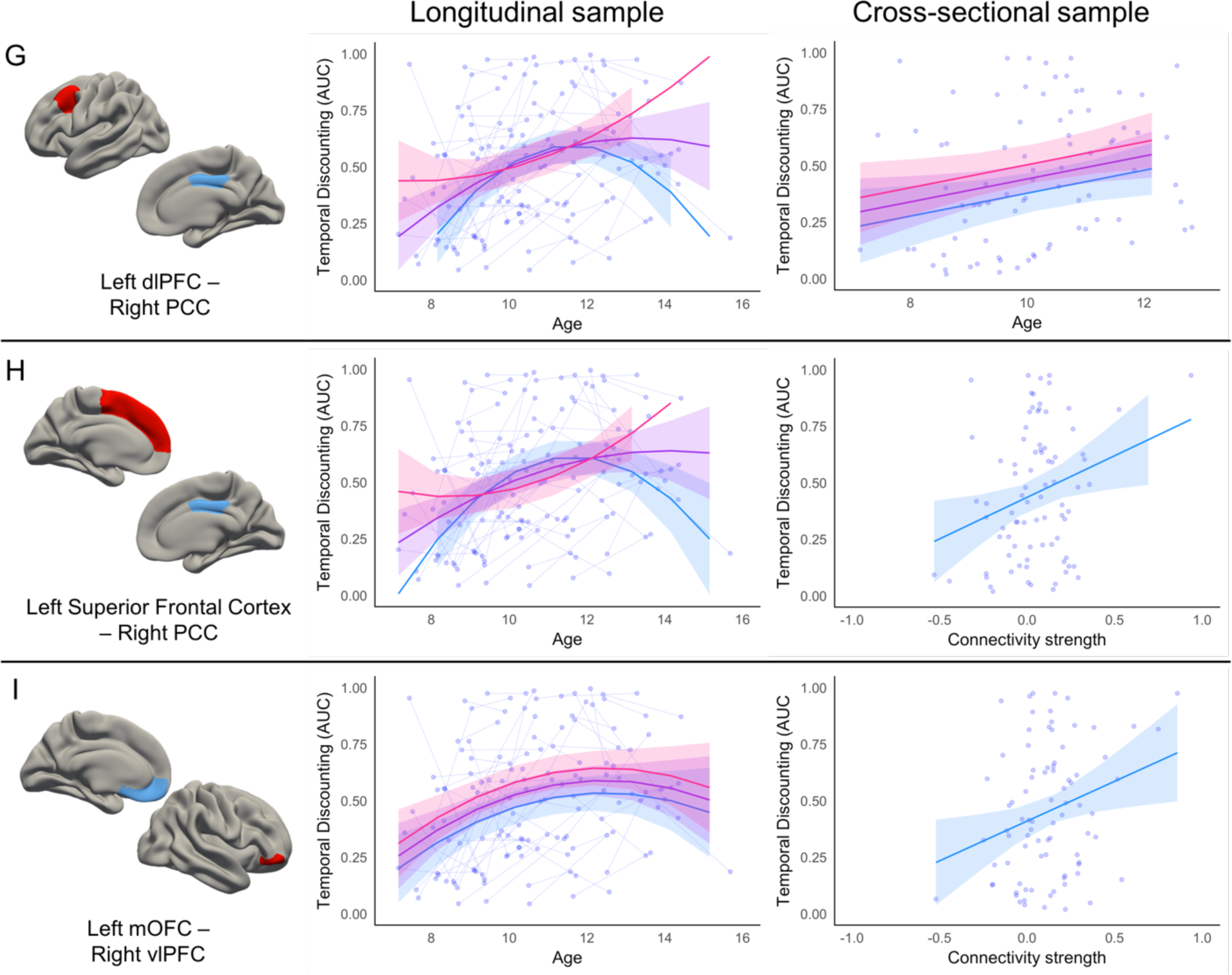
Relationship between valuation network and cognitive control network and AUC. The cortical regions involved in the connectivity between valuation system and cognitive control system are represented by blue and red, respectively, on the brain. The Pink trajectory represents AUC for an individual with 1 standard deviation higher connectivity than the mean between the two regions. Purple trajectory represents predicted AUC for participants with the mean connectivity strength between the two regions. Blue trajectory represents AUC for an individual with 1 standard deviation lower connectivity than the mean between the two regions. Raw data are plotted in the background, with each individual measurement representing a circle, and lines connecting data collected from the same individual across time.

### Role of dopaminergic signaling in temporal discounting behavior

All of the identified relevant connections between regions of the valuation network (amygdala, mOFC, PCC, and pallidum) showed a negative relationship with AUC, with stronger connectivity predicting a greater preference for SSR across participants. This could be related to the abundance of dopaminergic signaling in the valuation network. Multiple studies have shown that areas of the brain with dopaminergic innervation are involved in temporal discounting preference (Kobayashi & Schultz, 2008; Pine, Shiner, Seymour, & Dolan, 2010). Furthermore, it has been reported that individuals with increased dopamine release are more inclined to choose the SSR (Joutsa et al., 2015). Crossover work in animal models might allow for direct testing of this hypothesis (Grayson & Fair, 2017; Grayson, Kroenke, Neuringer, & Fair, 2014; Miranda-Dominguez et al., 2014; Stafford et al., 2014).

One hypothesis is that changes in the cortico-striatal circuitry that occur in the transition into adolescence are related to hormonal changes that affect the interaction within the networks (Blakemore, Burnett, & Dahl, 2010; Chambers, Taylor, & Potenza, 2003). Specifically, these hormonal changes impact and influence motivation towards reward seeking behaviors (Luciana & Collins, 2012). Pubertal hormones and neurotransmitters, such as sex hormones and dopamine, affect regions across the brain, but their effects (especially dopamine) on the vmPFC, NAcc, and caudate might influence the development of cognitive capacities such as abstract thinking, problem solving, and working memory (Chambers et al., 2003).

### Limitations and Future directions

This study examined temporal discounting preference as it relates to biological measures. However, social environmental factors can impact an individual’s subjective value of money and preference for waiting for a LLR. For example, a previous study found that individuals who grew up in lower socio-economic status environments (SES) preferred SSR, whereas individuals who grew up in higher SES environments preferred LLR (Griskevicius et al., 2013). In an experimental manipulation, Kidd, Palmeri, & Aslin (2013) demonstrated that children presented with a reliable environment demonstrated a significant increase in their delay time compared to children presented with an unstable environment. It should not be assumed that steeper discounting is always maladaptive. Very low socio-economic status populations were under-represented in the current study. Future investigations should assess how social environmental factors might impact the relationship between biological measures and temporal discounting preference.

Previous studies have shown evidence for heterogeneity in functional connectivity existing across individuals in typically developing as well as in clinical samples (Costa Dias et al., 2015; Fair, Bathula, Nikolas, & Nigg, 2012; Gates, Molenaar, Iyer, Nigg, & Fair, 2014). For example, graph theory and community detection can be used to classify typically developing children into specific neuropsychological subgroups (Gates et al., 2014), and functional subgroups can be differentiated based on heterogeneity related to behavioral characteristics including impulsivity (Costa Dias et al., 2015). This study did not account for these heterogeneity present in the group and further investigation should be considerate of this phenomenon. Further, the current study utilized a brain parcellation based on anatomical boundaries (the Desikan-Killiany atlas; Desikan et al., 2006) in order to test hypotheses generated from previous work. However, establishing the consistency of these findings with other parcellations (Glasser et al., 2016; Gordon et al., 2016) will be an important next step (Grayson & Fair, 2017; Hagmann, Grant, & Fair, 2012).

## Conclusion

On average, children start to prefer waiting for later, larger rewards as they transition into adolescence. However, there is a substantial amount of variability in temporal discounting preference between individuals across development. This study provides evidence that individual differences in functional brain connectivity within and between regions in cognitive control and valuation networks can account for variance in temporal discounting preference above age. Specifically, greater connectivity strength between cognitive control regions, as well as between cognitive control and valuation regions, was related to a preference for waiting for a larger reward. In contrast, greater connectivity strength between valuation network regions was related to a preference for taking an immediate, smaller, reward. Future studies should examine the impact of social environmental factors on the relationship between functional brain connectivity and temporal discounting behavior across development.

## Acknowledgements

We would like to thank the participants and families for their ongoing participation in this study, as well as the many staff involved in administrative tasks and data collection of this project.

## References

Achterberg, M., Peper, J. S., van Duijvenvoorde, A. C. K., Mandl, R. C. W., & Crone, E. A. (2016). Frontostriatal White Matter Integrity Predicts Development of Delay of Gratification: A Longitudinal Study. The Journal of Neuroscience: The Official Journal of the Society for Neuroscience, 36(6), 1954–1961. https://doi.org/10.1523/JNEUROSCI.3459-15.2016

Banich, M. T., De La Vega, A., Andrews-Hanna, J. R., Mackiewicz Seghete, K., Du, Y., & Claus, E. D. (2013). Developmental trends and individual differences in brain systems involved in intertemporal choice during adolescence. Psychology of Addictive Behaviors: Journal of the Society of Psychologists in Addictive Behaviors, 27(2), 416–430. https://doi.org/10.1037/a0031991

Blakemore, S.-J., Burnett, S., & Dahl, R. E. (2010). The role of puberty in the developing adolescent brain. Human Brain Mapping, 31(6), 926–933. https://doi.org/10.1002/hbm.21052

Boly, M., Phillips, C., Tshibanda, L., Vanhaudenhuyse, A., Schabus, M., Dang-Vu, T. T., … Laureys, S. (2008). Intrinsic brain activity in altered states of consciousness: how conscious is the default mode of brain function? Annals of the New York Academy of Sciences, 1129, 119–129. https://doi.org/10.1196/annals.1417.015

Burgess, G. C., Kandala, S., Nolan, D., Laumann, T. O., Power, J. D., Adeyemo, B., … Barch, D. M. (2016). Evaluation of Denoising Strategies to Address Motion-Correlated Artifacts in Resting-State Functional Magnetic Resonance Imaging Data from the Human Connectome Project. Brain Connectivity, 6(9), 669–680. https://doi.org/10.1089/brain.2016.0435

Calluso, C., Tosoni, A., Pezzulo, G., Spadone, S., & Committeri, G. (2015). Interindividual Variability in Functional Connectivity as Long-Term Correlate of Temporal Discounting. PLOS ONE, 10(3), e0119710. https://doi.org/10.1371/journal.pone.0119710

Casey, B. J. (2015). Beyond simple models of self-control to circuit-based accounts of adolescent behavior. Annual Review of Psychology, 66, 295–319. https://doi.org/10.1146/annurev-psych-010814-015156

Chambers, R. A., Taylor, J. R., & Potenza, M. N. (2003). Developmental neurocircuitry of motivation in adolescence: a critical period of addiction vulnerability. The American Journal of Psychiatry, 160(6), 1041–1052.

Christakou, A., Brammer, M., & Rubia, K. (2011). Maturation of limbic corticostriatal activation and connectivity associated with developmental changes in temporal discounting. NeuroImage, 54(2), 1344–1354. https://doi.org/10.1016/j.neuroimage.2010.08.067

Conners, C. K. (2003). Conners’ rating scales: Revised technical manual. New York, NY: Multi-Health Systems. Retrieved from https://www.mhs.com/MHS-Assessment?prodname=conners3

Costa Dias, T. G., Iyer, S. P., Carpenter, S. D., Cary, R. P., Wilson, V. B., Mitchell, S. H., … Fair, D. A. (2015). Characterizing heterogeneity in children with and without ADHD based on reward system connectivity. Developmental Cognitive Neuroscience, 11, 155–174. https://doi.org/10.1016/j.dcn.2014.12.005

Costa Dias, T. G., Wilson, V. B., Bathula, D. R., Iyer, S. P., Mills, K. L., Thurlow, B. L., … Fair, D. A. (2012). Reward circuit connectivity relates to delay discounting in children with attention-deficit/hyperactivity disorder. European Neuropsychopharmacology: The Journal of the European College of Neuropsychopharmacology. https://doi.org/10.1016/j.euroneuro.2012.10.015

Cross, C. P., Copping, L. T., & Campbell, A. (2011). Sex differences in impulsivity: a meta-analysis. Psychological Bulletin, 137(1), 97–130. https://doi.org/10.1037/a0021591

Dale, A. M., Fischl, B., & Sereno, M. I. (1999). Cortical Surface-Based Analysis: I. Segmentation and Surface Reconstruction. NeuroImage, 9(2), 179–194. https://doi.org/10.1006/nimg.1998.0395

de Water, E., Cillessen, A. H. N., & Scheres, A. (2014). Distinct age-related differences in temporal discounting and risk taking in adolescents and young adults. Child Development, 85(5), 1881–1897. https://doi.org/10.1111/cdev.12245

Desikan, R. S., Ségonne, F., Fischl, B., Quinn, B. T., Dickerson, B. C., Blacker, D., … Killiany, R. J. (2006). An automated labeling system for subdividing the human cerebral cortex on MRI scans into gyral based regions of interest. NeuroImage, 31(3), 968–980. https://doi.org/10.1016/j.neuroimage.2006.01.021

Dosenbach, N. U. F., Koller, J. M., Earl, E. A., Miranda-Dominguez, O., Klein, R. L., Van, A. N., … Fair, D. A. (2017). Real-time motion analytics during brain MRI improve data quality and reduce costs. NeuroImage, 161, 80–93. https://doi.org/10.1016/j.neuroimage.2017.08.025

Fair, D. A., Bathula, D., Nikolas, M. A., & Nigg, J. T. (2012). Distinct neuropsychological subgroups in typically developing youth inform heterogeneity in children with ADHD. Proceedings of the National Academy of Sciences of the United States of America, 109(17), 6769–6774. https://doi.org/10.1073/pnas.1115365109

Fair, D. A., Nigg, J. T., Iyer, S., Bathula, D., Mills, K. L., Dosenbach, N. U. F., … Milham, M. P. (2012). Distinct neural signatures detected for ADHD subtypes after controlling for micro-movements in resting state functional connectivity MRI data. Frontiers in Systems Neuroscience, 6, 80. https://doi.org/10.3389/fnsys.2012.00080

Fischl, B., & Dale, A. M. (2000). Measuring the thickness of the human cerebral cortex from magnetic resonance images. Proceedings of the National Academy of Sciences, 97(20), 11050–11055. https://doi.org/10.1073/pnas.200033797

Fischl, B., Salat, D. H., Busa, E., Albert, M., Dieterich, M., Haselgrove, C., … Dale, A. M. (2002). Whole brain segmentation: automated labeling of neuroanatomical structures in the human brain. Neuron, 33(3), 341–355.

Fischl, B., Sereno, M. I., & Dale, A. M. (1999). Cortical surface-based analysis. II: Inflation, flattening, and a surface-based coordinate system. NeuroImage, 9(2), 195–207. https://doi.org/10.1006/nimg.1998.0396

Fischl, B., van der Kouwe, A., Destrieux, C., Halgren, E., Ségonne, F., Salat, D. H., … Dale, A. M. (2004). Automatically parcellating the human cerebral cortex. Cerebral Cortex (New York, N.Y.: 1991), 14(1), 11–22.

Gates, K. M., Molenaar, P. C. M., Iyer, S. P., Nigg, J. T., & Fair, D. A. (2014). Organizing heterogeneous samples using community detection of GIMME-derived resting state functional networks. PloS One, 9(3), e91322. https://doi.org/10.1371/journal.pone.0091322

Glasser, M. F., Coalson, T. S., Robinson, E. C., Hacker, C. D., Harwell, J., Yacoub, E., … Van Essen, D. C. (2016). A multi-modal parcellation of human cerebral cortex. Nature, 536(7615), 171–178. https://doi.org/10.1038/nature18933

Glasser, M. F., Sotiropoulos, S. N., Wilson, J. A., Coalson, T. S., Fischl, B., Andersson, J. L., … Jenkinson, M. (2013). The minimal preprocessing pipelines for the Human Connectome Project. NeuroImage, 80(Supplement C), 105–124. https://doi.org/10.1016/j.neuroimage.2013.04.127

Gordon, E. M., Laumann, T. O., Adeyemo, B., Huckins, J. F., Kelley, W. M., & Petersen, S. E. (2016). Generation and Evaluation of a Cortical Area Parcellation from Resting-State Correlations. Cerebral Cortex (New York, N.Y.: 1991), 26(1), 288–303. https://doi.org/10.1093/cercor/bhu239

Grayson, D. S., & Fair, D. A. (2017). Development of large-scale functional networks from birth to adulthood: A guide to the neuroimaging literature. NeuroImage, 160, 15–31. https://doi.org/10.1016/j.neuroimage.2017.01.079

Grayson, D. S., Kroenke, C. D., Neuringer, M., & Fair, D. A. (2014). Dietary omega-3 fatty acids modulate large-scale systems organization in the rhesus macaque brain. The Journal of Neuroscience: The Official Journal of the Society for Neuroscience, 34(6), 2065–2074. https://doi.org/10.1523/JNEUROSCI.3038-13.2014

Green, L., Myerson, J., & Mcfadden, E. (1997). Rate of temporal discounting decreases with amount of reward. Memory & Cognition, 25(5), 715–723. https://doi.org/10.3758/BF03211314

Greve, D. N., & Fischl, B. (2009). Accurate and robust brain image alignment using boundary-based registration. NeuroImage, 48(1), 63–72. https://doi.org/10.1016/j.neuroimage.2009.06.060

Griskevicius, V., Ackerman, J. M., Cantú, S. M., Delton, A. W., Robertson, T. E., Simpson, J. A., … Tybur, J. M. (2013). When the economy falters, do people spend or save? Responses to resource scarcity depend on childhood environments. Psychological Science, 24(2), 197–205. https://doi.org/10.1177/0956797612451471

Haber, S. N., & Knutson, B. (2009). The Reward Circuit: Linking Primate Anatomy and Human Imaging. Neuropsychopharmacology, 35(1), 4–26. https://doi.org/10.1038/npp.2009.129

Hagmann, P., Grant, P. E., & Fair, D. A. (2012). MR connectomics: a conceptual framework for studying the developing brain. Frontiers in Systems Neuroscience, 6, 43. https://doi.org/10.3389/fnsys.2012.00043

Hamilton, K. R., Mitchell, M. R., Wing, V. C., Balodis, I. M., Bickel, W. K., Fillmore, M., … Moeller, F. G. (2015). Choice impulsivity: Definitions, measurement issues, and clinical implications. Personality Disorders, 6(2), 182–198. https://doi.org/10.1037/per0000099

Hare, T. A., Camerer, C. F., & Rangel, A. (2009). Self-Control in Decision-Making Involves Modulation of the vmPFC Valuation System. Science, 324(5927), 646–648. https://doi.org/10.1126/science.1168450

Hare, T. A., Hakimi, S., & Rangel, A. (2014). Activity in dlPFC and its effective connectivity to vmPFC are associated with temporal discounting. Frontiers in Neuroscience, 8. https://doi.org/10.3389/fnins.2014.00050

Jenkinson, M., Beckmann, C. F., Behrens, T. E. J., Woolrich, M. W., & Smith, S. M. (2012). FSL. NeuroImage, 62(2), 782–790. https://doi.org/10.1016/j.neuroimage.2011.09.015

Johnson, M. W., & Bickel, W. K. (2008). An algorithm for identifying nonsystematic delay discounting data. Experimental and Clinical Psychopharmacology, 16(3), 264–274. https://doi.org/10.1037/1064-1297.16.3.264

Joutsa, J., Voon, V., Johansson, J., Niemelä, S., Bergman, J., & Kaasinen, V. (2015). Dopaminergic function and intertemporal choice. Translational Psychiatry, 5, e491. https://doi.org/10.1038/tp.2014.133

Karalunas, S. L., Gustafsson, H. C., Dieckmann, N. F., Tipsord, J., Mitchell, S. H., & Nigg, J. T. (2017). Heterogeneity in development of aspects of working memory predicts longitudinal attention deficit hyperactivity disorder symptom change. Journal of Abnormal Psychology, 126(6), 774–792. https://doi.org/10.1037/abn0000292

Kidd, C., Palmeri, H., & Aslin, R. N. (2013). Rational snacking: young children’s decision making on the marshmallow task is moderated by beliefs about environmental reliability. Cognition, 126(1), 109–114. https://doi.org/10.1016/j.cognition.2012.08.004

Kobayashi, S., & Schultz, W. (2008). Influence of reward delays on responses of dopamine neurons. The Journal of Neuroscience: The Official Journal of the Society for Neuroscience, 28(31), 7837–7846. https://doi.org/10.1523/JNEUROSCI.1600-08.2008

Lee, N. C., de Groot, R. H. M., Boschloo, A., Dekker, S., Krabbendam, L., & Jolles, J. (2013). Age and educational track influence adolescent discounting of delayed rewards. Frontiers in Psychology, 4. https://doi.org/10.3389/fpsyg.2013.00993

Li, N., Ma, N., Liu, Y., He, X.-S., Sun, D.-L., Fu, X.-M., … Zhang, D.-R. (2013). Resting-state functional connectivity predicts impulsivity in economic decision-making. The Journal of Neuroscience: The Official Journal of the Society for Neuroscience, 33(11), 4886–4895. https://doi.org/10.1523/JNEUROSCI.1342-12.2013

Luciana, M., & Collins, P. F. (2012). Incentive Motivation, Cognitive Control, and the Adolescent Brain: Is It Time for a Paradigm Shift? Child Development Perspectives, 6(4), 392–399. https://doi.org/10.1111/j.1750-8606.2012.00252.x

Miranda-Dominguez, O., Mills, B. D., Grayson, D., Woodall, A., Grant, K. A., Kroenke, C. D., & Fair, D. A. (2014). Bridging the gap between the human and macaque connectome: a quantitative comparison of global interspecies structure-function relationships and network topology. The Journal of Neuroscience: The Official Journal of the Society for Neuroscience, 34(16), 5552–5563. https://doi.org/10.1523/JNEUROSCI.4229-13.2014

Mitchell, S. H., Wilson, V. B., & Karalunas, S. L. (2015). Comparing hyperbolic, delay-amount sensitivity and present-bias models of delay discounting. Behavioural Processes, 114, 52–62. https://doi.org/10.1016/j.beproc.2015.03.006

Myerson, J., & Green, L. (1995). Discounting of delayed rewards: Models of individual choice. Journal of the Experimental Analysis of Behavior, 64(3), 263–276. https://doi.org/10.1901/jeab.1995.64-263

Myerson, J., Green, L., & Warusawitharana, M. (2001). Area under the curve as a measure of discounting. Journal of the Experimental Analysis of Behavior, 76(2), 235–243. https://doi.org/10.1901/jeab.2001.76-235

Odum, fA. L. (2011). Delay Discounting: I’m a k, You’re a k. Journal of the Experimental Analysis of Behavior, 96(3), 427–439. https://doi.org/10.1901/jeab.2011.96-423

Olson, E. A., Collins, P. F., Hooper, C. J., Muetzel, R., Lim, K. O., & Luciana, M. (2009). White Matter Integrity Predicts Delay Discounting Behavior in 9- to 23-Year-Olds: A Diffusion Tensor Imaging Study. Journal of Cognitive Neuroscience, 21(7), 1406–1421. https://doi.org/10.1162/jocn.2009.21107

Peters, J., & Büchel, C. (2010). Episodic future thinking reduces reward delay discounting through an enhancement of prefrontal-mediotemporal interactions. Neuron, 66(1), 138–148. https://doi.org/10.1016/j.neuron.2010.03.026

Peters, J., & Büchel, C. (2011). The neural mechanisms of inter-temporal decision-making: understanding variability. Trends in Cognitive Sciences, 15(5), 227–239. https://doi.org/10.1016/j.tics.2011.03.002

Pine, A., Shiner, T., Seymour, B., & Dolan, R. J. (2010). Dopamine, Time, and Impulsivity in Humans. Journal of Neuroscience, 30(26), 8888–8896. https://doi.org/10.1523/JNEUROSCI.6028-09.2010

Power, J. D., Barnes, K. A., Snyder, A. Z., Schlaggar, B. L., & Petersen, S. E. (2012). Spurious but systematic correlations in functional connectivity MRI networks arise from subject motion. NeuroImage, 59(3), 2142–2154. https://doi.org/10.1016/j.neuroimage.2011.10.018

Power, J. D., Mitra, A., Laumann, T. O., Snyder, A. Z., Schlaggar, B. L., & Petersen, S. E. (2014). Methods to detect, characterize, and remove motion artifact in resting state fMRI. NeuroImage, 84, 320–341. https://doi.org/10.1016/j.neuroimage.2013.08.048

Power, J. D., Schlaggar, B. L., & Petersen, S. E. (2014). Studying Brain Organization via Spontaneous fMRI Signal. Neuron, 84(4), 681–696. https://doi.org/10.1016/j.neuron.2014.09.007

Puig-Antich, J., & Ryan, N. (1986). Schedule for Affective Disorders and Schizophrenia for School-Age Children. Pittsburgh, PA: Western Psychiatric Institute and Clinic.

Richards, J. B., Zhang, L., Mitchell, S. H., & de Wit, H. (1999). Delay or probability discounting in a model of impulsive behavior: effect of alcohol. Journal of the Experimental Analysis of Behavior, 71(2), 121–143. https://doi.org/10.1901/jeab.1999.71-121

Sadaghiani, S., & Kleinschmidt, A. (2013). Functional interactions between intrinsic brain activity and behavior. NeuroImage, 80, 379–386. https://doi.org/10.1016/j.neuroimage.2013.04.100

Scheres, A., de Water, E., & Mies, G. W. (2013). The neural correlates of temporal reward discounting. Wiley Interdisciplinary Reviews. Cognitive Science, 4(5), 523–545. https://doi.org/10.1002/wcs.1246

Scheres, A., Tontsch, C., Thoeny, A. L., & Sumiya, M. (2014). Temporal reward discounting in children, adolescents, and emerging adults during an experiential task. Developmental Psychology, 5, 711. https://doi.org/10.3389/fpsyg.2014.00711

Smith, S. M., Jenkinson, M., Woolrich, M. W., Beckmann, C. F., Behrens, T. E. J., Johansen-Berg, H., … Matthews, P. M. (2004). Advances in functional and structural MR image analysis and implementation as FSL. NeuroImage, 23 Suppl 1, S208-219. https://doi.org/10.1016/j.neuroimage.2004.07.051

Stafford, J. M., Jarrett, B. R., Miranda-Dominguez, O., Mills, B. D., Cain, N., Mihalas, S., … Fair, D. A. (2014). Large-scale topology and the default mode network in the mouse connectome. Proceedings of the National Academy of Sciences of the United States of America, 111(52), 18745–18750. https://doi.org/10.1073/pnas.1404346111

Steinberg, L., Graham, S., O’Brien, L., Woolard, J., Cauffman, E., & Banich, M. (2009). Age differences in future orientation and delay discounting. Child Development, 80(1), 28–44. https://doi.org/10.1111/j.1467-8624.2008.01244.x

van den Bos, W., & McClure, S. M. (2013). Towards a general model of temporal discounting. Journal of the Experimental Analysis of Behavior, 99(1), 58–73. https://doi.org/10.1002/jeab.6

van den Bos, W., Rodriguez, C. A., Schweitzer, J. B., & McClure, S. M. (2014). Connectivity Strength of Dissociable Striatal Tracts Predict Individual Differences in Temporal Discounting. The Journal of Neuroscience, 34(31), 10298–10310. https://doi.org/10.1523/JNEUROSCI.4105-13.2014

van den Bos, W., Rodriguez, C. A., Schweitzer, J. B., & McClure, S. M. (2015). Adolescent impatience decreases with increased frontostriatal connectivity. Proceedings of the National Academy of Sciences, 112(29), E3765–E3774. https://doi.org/10.1073/pnas.1423095112

Volkow, N. D., & Baler, R. D. (2015). NOW vs LATER brain circuits: implications for obesity and addiction. Trends in Neurosciences, 38(6), 345–352. https://doi.org/10.1016/j.tins.2015.04.002

Wang, S., Zhou, M., Chen, T., Yang, X., Chen, G., & Gong, Q. (2017). Delay discounting is associated with the fractional amplitude of low-frequency fluctuations and resting-state functional connectivity in late adolescence. Scientific Reports, 7(1), 10276. https://doi.org/10.1038/s41598-017-11109-z

Wechsler, D. (2003). Wechsler Intelligence Scale for Children, 4th ed., (WISC-IV) technical and interpretive manual. San Antonio, TX: Harcourt Brace.

Woolrich, M. W., Jbabdi, S., Patenaude, B., Chappell, M., Makni, S., Behrens, T., … Smith, S. M. (2009). Bayesian analysis of neuroimaging data in FSL. NeuroImage, 45(1 Suppl), S173-186. https://doi.org/10.1016/j.neuroimage.2008.10.055

Yi, R., Pitcock, J. A., Landes, R. D., & Bickel, W. K. (2010). The short of it: abbreviating the temporal discounting procedure. Experimental and Clinical Psychopharmacology, 18(4), 366–374. https://doi.org/10.1037/a0019904

